# Lysine 117 on ataxin-3 modulates toxicity in *Drosophila* models of Spinocerebellar Ataxia Type 3

**DOI:** 10.1101/2023.05.30.542896

**Authors:** Jessica R. Blount, Nikhil C. Patel, Kozeta Libohova, Autumn L. Harris, Wei-Ling Tsou, Alyson Sujkowski, Sokol V. Todi

**Affiliations:** Department of Pharmacology, Wayne State University; Maximizing Access to Research Careers, Wayne State University; Department of Neurology, Wayne State University

## Abstract

Ataxin-3 (Atxn3) is a deubiquitinase with a polyglutamine (polyQ) repeat tract whose abnormal expansion causes the neurodegenerative disease, Spinocerebellar Ataxia Type 3 (SCA3; also known as Machado-Joseph Disease). The ubiquitin chain cleavage properties of Atxn3 are enhanced when it is ubiquitinated at lysine (K) at position 117. K117-ubiqutinated Atxn3 cleaves poly-ubiquitin more rapidly in vitro compared to its unmodified counterpart and this residue is also important for Atxn3 roles in cell culture and in *Drosophila melanogaster*. How polyQ expansion causes SCA3 remains unclear. To gather insight into the biology of disease of SCA3, here we posited the question: is K117 important for toxicity caused by Atxn3? We generated transgenic *Drosophila* lines that express full-length, human, pathogenic Atxn3 with 80 polyQ with an intact or mutated K117. We found that K117 mutation mildly enhances the toxicity and aggregation of pathogenic Atxn3 in *Drosophila*. An additional transgenic line that expresses Atxn3 without any K residues confirms increased aggregation of pathogenic Atxn3 whose ubiquitination is perturbed. These findings suggest Atxn3 ubiquitination as a regulatory step of SCA3, in part by modulating its aggregation.

## INTRODUCTION

The most common, dominantly inherited ataxia worldwide, Spinocerebellar Ataxia Type 3 (SCA3), also known as Machado-Joseph Disease, is caused by a mutation in the deubiquitinase (DUB) ataxin-3 (Atxn3) (Kawaguchi et al., 1994; Stevanin et al., 1995a; Stevanin et al., 1995b). This mutation is an abnormal expansion in a CAG triplet repeat in the gene, *ATXN3*, translated into polyglutamine (polyQ) (Johnson et al., 2022b; Paulson et al., 2017; Zoghbi and Orr, 2009). Nine different genes have similar repeats, whose abnormal lengthening causes polyQ expansion in proteins and causes inherited neurodegenerative disorders that collectively comprise the polyQ family of diseases (Johnson et al., 2022b; Paulson et al., 2017; Zoghbi and Orr, 2009).

It is unclear how polyQ expansion in Atxn3 causes SCA3. According to data from animal and cell models of SCA3, polyQ expansion may modify normal functions of Atxn3 as well as endow it with new properties (Dantuma and Herzog, 2020; Johnson et al., 2019; Johnson et al., 2021; Johnson et al., 2022a; Johnson et al., 2020; Matos et al., 2019a; Ristic et al., 2018; Sutton et al., 2017; Todi et al., 2007). Still, precise molecular sequelae remain uncertain and would benefit from systematic investigations of the disease protein, its partners, and its functions. Towards that end, we have explored the role of ubiquitin (its substrate) on Atxn3 (Faggiano et al., 2013; Faggiano et al., 2015; Todi et al., 2010; Todi et al., 2009; Winborn et al., 2008). We found that: wild-type and pathogenic Atxn3 are prominently ubiquitinated in vitro, in cultured cells, in *Drosophila*, and in mouse brain; wild-type and pathogenic Atxn3 are primarily ubiquitinated at lysine (K) at position 117, which resides on the catalytic domain (figure 1A); the in vitro catalytic activity of wild-type and pathogenic Atxn3 is markedly enhanced by its own ubiquitination; neither wild-type, nor pathogenic Atxn3 require their own ubiquitination to be degraded by the proteasome; and ubiquitination of Atxn3 at K117 is important for functions of Atxn3 in cell culture and in *Drosophila* (Blount et al., 2020; Blount et al., 2014; Faggiano et al., 2013; Faggiano et al., 2015; Todi et al., 2010; Todi et al., 2009; Tsou et al., 2013). These prior results led us to ask: is ubiquitination of Atxn3 important for its toxicity? We reasoned that this question would bear special importance to further understand the pathogenic properties of the SCA3 protein, providing additional pieces to the larger puzzle of the SCA3 biology of disease and the need for therapeutics to treat it.

**Figure 1:**
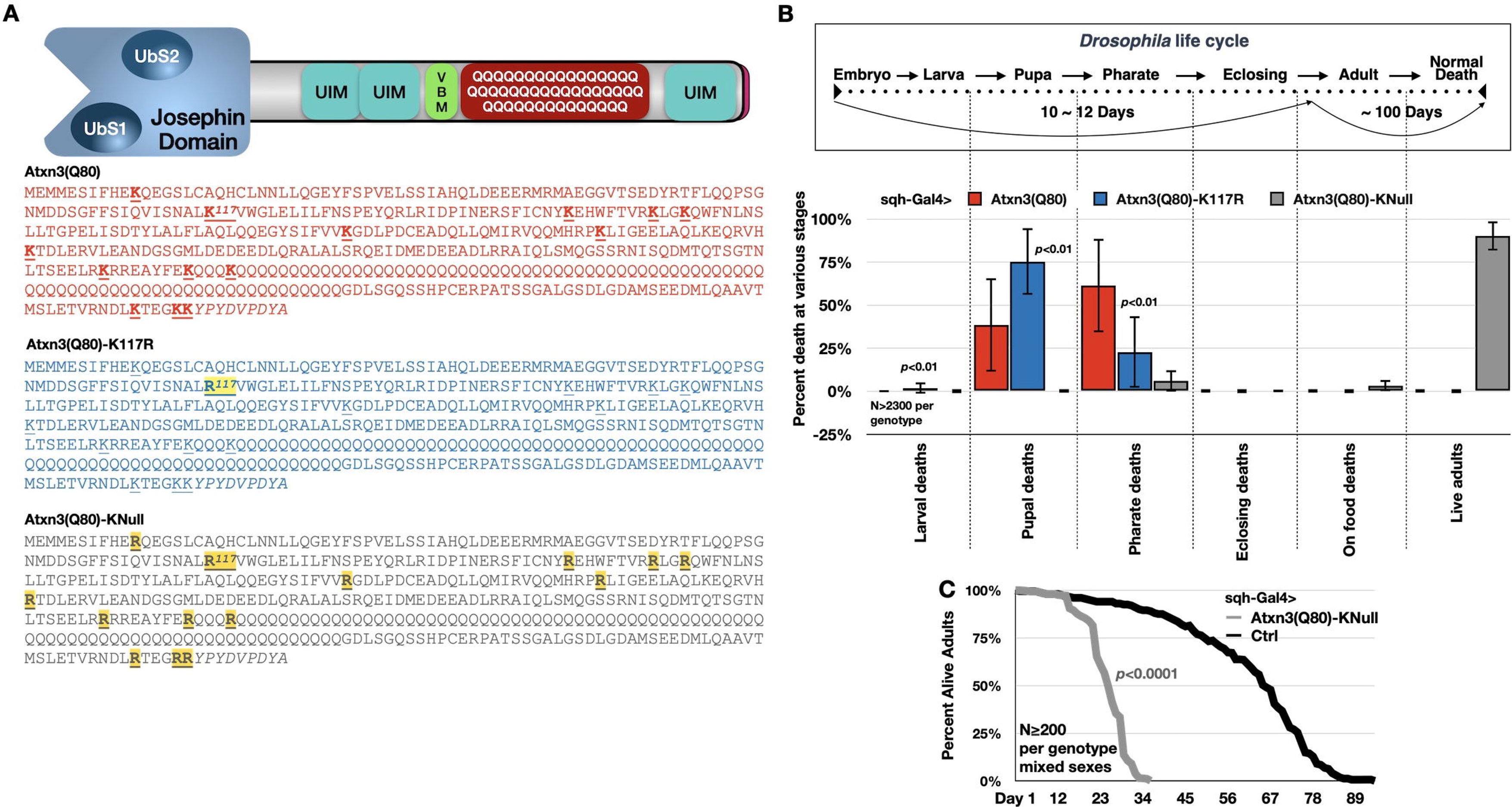
K117 mutation enhances Atxn3 toxicity in all tissues. A) Atxn3 protein domains and the sequences encoded by the transgenes in this study. UbS: ubiquitin-binding site. UIM: ubiquitin-binding motif. VBM: VCP-binding motif. HA tag is appended inline to each protein (sequence in italics). Josephin domain is the catalytic portion of Atxn3. B) Results of sqh-Gal4 (ubiquitous driver) expressing the indicated Atxn3 forms. Ctrl: sqh-Gal4 in trans with the parental line used to generate the Atrxn3 transgenics. Top: mean -/+ SD. Statistics: t-tests comparing Atxn3(Q80)-K117R to Atxn3(Q80). Bottom: statistics: log-rank test.

Here, we report the generation and examination of additional *Drosophila* models of SCA3 that utilize the binary Gal4-UAS expression system either constitutively or in a spatially and temporally restricted manner. These transgenic lines express human Atxn3 protein with an expanded polyQ repeat with intact or mutated K117, or absent any lysine residues. We found that K117 mutation enhances the toxicity of pathogenic Atxn3 in the fruit fly, and that this observation coincides with mildly increased aggregation of the offending protein. Overall, our studies point to a modest, but significant modifier role for ubiquitination of Atxn3 at K117 in its toxicity.

## RESULTS

### Mutation of K117 enhances toxicity of pathogenic Atxn3 in *Drosophila*

To examine the role of K117 in the toxicity of pathogenic Atxn3 (figure 1A), we generated *Drosophila melanogaster* transgenic lines that express one of the following versions of the SCA3 protein: Atxn3 with 80Q (within human disease range) without any additional mutations; Atxn3 with 80Q where K117 is mutated into the similar, but not ubiquitinatable, amino acid arginine (K117R); and a version with 80Q but where each of the lysine residues is mutated into arginines (KNull; figure 1A). The transgenes consist of the full-length, human cDNA. Each of the transgenes is C-terminally tagged with an HA epitope. The KNull line was generated to inquire into the role of ubiquitination more generally on Atxn3 toxicity, since we have shown before that while K117 is the primary site of ubiquitination, other lysines on Atxn3 can also be modified (Todi et al., 2010).

We started by examining the toxicity of each transgene when expressed in all tissues, at all stages. Driven by sqh-Gal4, we noticed that expression of Atxn3(Q80) or K117R both led to lethality during fly development as pupae and pharate adults (figure 1B). Statistically, K117R led to a higher percentage of developing flies dying as pupae rather than pharate adults, compared to Atxn3(Q80). In neither case did we observe any adult flies eclose successfully from their pupal cases (figure 1B). KNull expression, however, significantly improved developmental lethality. Although most KNull flies emerged as adults and survived for ∼30 days, longevity was far shorter than the longevity of flies not expressing pathogenic Atxn3 (figure 1C). We conclude that lysine mutations impact the toxicity of pathogenic Atxn3 when expressed in all tissues in the fly.

Next, we turned our attention to the impact of pathogenic Atxn3 expression in neurons, the tissue most impacted in SCA3. We expressed each version of pathogenic Atxn3 in all neurons throughout the fly life, or only in adults (figure 2). As shown in figure 2A, constitutive neuronal expression led to differential toxicity: Atxn3(Q80) was more toxic than KNull, but less so than K117R in males and females. Similarly, expression of these transgenes only in adult fly neurons led to similar differences in toxicity. These results indicate that mutation of K117 enhances the toxicity of pathogenic Atxn3, whereas mutation of all lysine residues reduces it.

**Figure 2:**
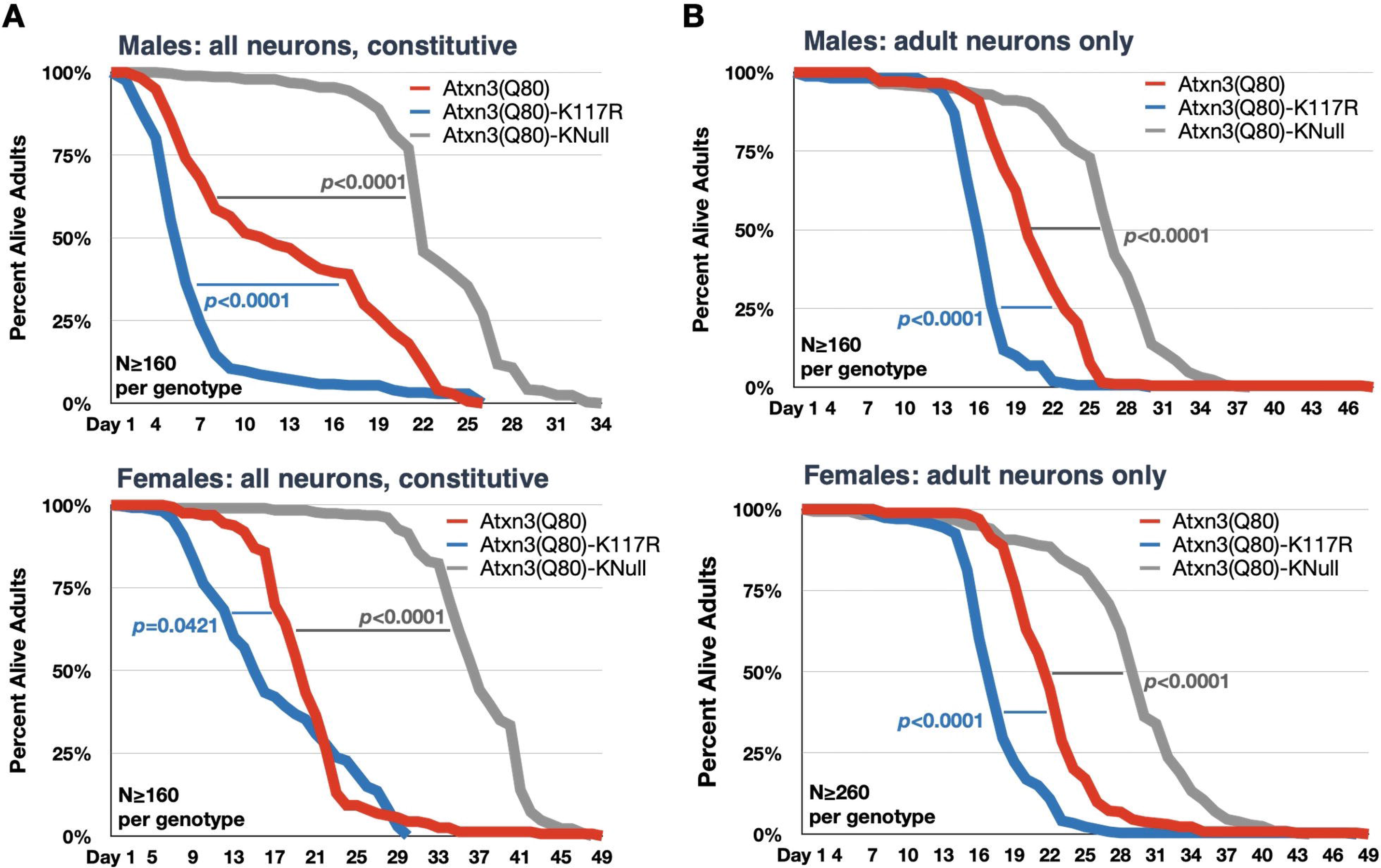
K117 mutation enhances Atxn3 toxicity in neurons. A, B) Longevity outcomes for adult flies expressing the noted versions of Atxn3 in all neuronal cells. Statistics: log-rank tests.

### Mutation of K117 does not impact the turnover or distribution of pathogenic Atxn3 in flies

We then turned our attention to biochemical properties of the Atxn3 versions expressed above. Constitutive, pan-neuronal expression of Atxn3(Q80), K117R and KNull led to similar overall protein levels for Atxn3(Q80) and K117R, but markedly lower levels for KNull (figure 3A).

**Figure 3:**
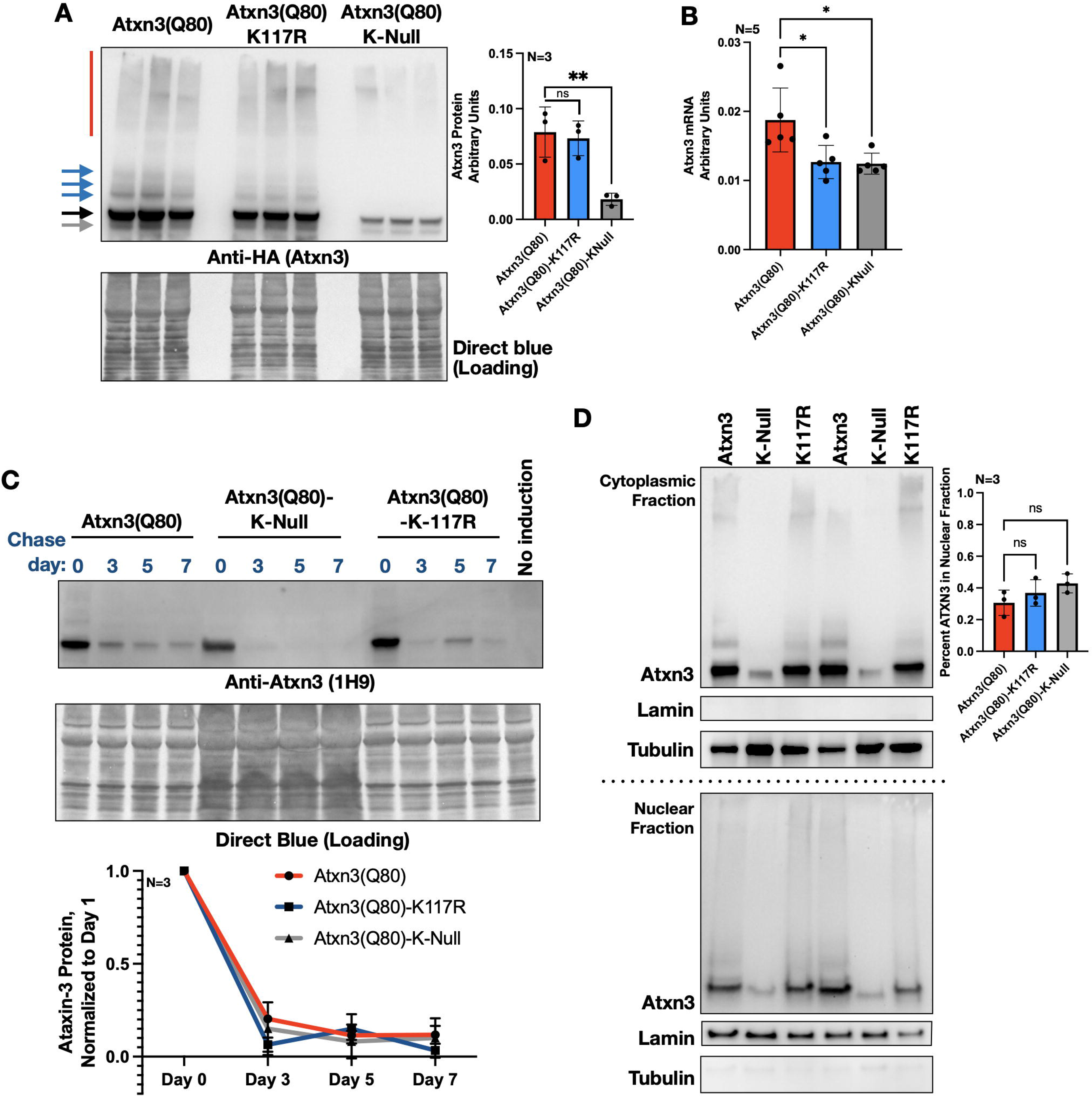
Lysine mutations do not impact turnover or subcellular fractionation of pathogenic Atxn3. A) Western blots from adult female flies expressing the noted versions of Atxn3 in all neurons, constitutively. Day 1 adults. Each lane represents an independent repeat. Black arrow: main, unmodified Atxn3 band. Blue arrows: ubiquitinated Atxn3. Gray arrow: likely proteolytic fragment we observe sometimes. Red bar: SDS-resistant Atxn3. Quantifications: images from the left were used for quantification, normalized to their respective Direct blue loading control signal. The entire Atxn3 signal was utilized, main band to top. Means -/+ SD. Statistics: one-way ANOVA with Dunnett’s post hoc. **: p<0.001 B) qRT-PCR results from 1-day-old adult females constitutively expressing the noted Atxn3 versions in all neurons. Means -/+ SD. Statistics: one-way ANOVA with Dunnett’s post hoc. *: p<0.05. C) Top: Western blots from adult female flies expressing the noted versions of Atxn3 in neurons for 7 days (+RU486) then allowed to degrade Atxn3 over the indicated timeline (-RU486). A higher amount of lysates from KNull was loaded to equilibrate as much as possible starting amounts at day 0. Bottom: quantification of signal from the top and additional independent repeats, normalized to respective day 0 levels. No statistical significances found with one-way ANOVA with Dunnett’s post hoc among day 3, 5 and 7 comparisons. D) Left: Western blots of nuclear/cytoplasmic fractionations of Atxn3 versions noted expressed constitutively in all neurons. Flies were 1-day-old females. Right: quantification of images from the left and other independent repeats. Means -/+ SD. Statistics: one-way ANOVA with Dunnett’s post hoc. ns: not significant.

According to qRT-PCR, the mRNA levels of Atxn3(Q80) were higher than those of K117R and KNull, the latter two being comparable (figure 3B). We had previously observed that lysine-to-arginine mutations impact the mRNA levels of wild-type Atxn3 expressed in cultured mammalian cells in the opposite direction of what we noticed here: Atxn3(Q12) mRNA levels are higher when nucleotide mutations substitute all lysines for arginines (Blount et al., 2020). This difference in findings may be due to differential handling of Atxn3 mRNA in culture versus in vivo, and could be further due to differences in CAG repeats, 80 in this case and 12 in the prior study.

To examine whether lysine mutations impact the turnover of Atxn3 protein in fly neurons, we induced their expression in adult neurons for 7 days, and then removed the inducer (RU486) to allow the accumulated protein to dissipate. As summarized in figure 3C, the rate of dissipation of the three versions of Atxn3 protein is similar, regardless of which lysine residues are present. We interpret these data to indicate that overall ubiquitination of Atxn3 – or only at K117 – is not necessary for its turnover (we have reported this before in mammalian cells and with non-pathogenic Atxn3 in *Drosophila* (Blount et al., 2020; Blount et al., 2014; Todi et al., 2010)). According to these results, differences in toxicity between Atxn3(Q80) and K117R are unlikely to be due to difference in the overall protein levels; however, reduced toxicity from KNull likely reflects the overall lower protein levels of this construct. We presume that the lower levels of KNull are due to a combination of reduced mRNA levels and potentially translation-dependent degradation. The fact that differences in the mRNA levels between Atxn3(Q80) and K117R do not translate into differences at the overall protein level may be due to differences in the rates of translation versus mRNA stability between these two constructs; we did not pursue these possibilities further in this work.

Nuclear localization of pathogenic Atxn3 is particularly problematic in mouse models of SCA3 (Bichelmeier et al., 2007; Schmidt et al., 1998). We examined the sub-cellular fractionation of Atxn3 biochemically. We observed a mild trend of increased nucellar presence when K117 or all lysines are mutated, but this did not reach statistical significance (figure 3D), suggesting that differences in toxicity between Atxn3(Q80) and K117R are unlikely to stem from differences in sub-cellular localization.

### DnaJ-1 suppresses the toxicity of pathogenic Atxn3 independently of K117

We and other reported that exogenous expression of various folding-related proteins has marked impact on the toxicity of pathogenic Atxn3 in *Drosophila* and other models of SCA3 (Johnson et al., 2021; Johnson et al., 2020; Joshi et al., 2021; Kakkar et al., 2012; Kazemi-Esfarjani and Benzer, 2000; Kuhlbrodt et al., 2011; Ristic et al., 2018; Tsou et al., 2015a; Tsou et al., 2015b). We wondered whether the protective effect of one such protein, DnaJ-1, which interacts with Atxn3 and is regulated by it depends on lysine residues of the SCA3 protein: could the differences in toxicity between Atxn3(Q80) and K117R be due to their differential response to cellular attempts to reduce their toxicity?

We expressed either of the three versions of pathogenic Atxn3 in adult fly neurons in the absence or presence of exogenous DnaJ-1. We found that in each case, exogenous DnaJ-1 led to improved longevity regardless of sex or lysine mutations (figure 4A, B). Co- immunopurifications (co-IPs) from the same flies did not reveal significant differences in the ability of Atxn3(Q80), K117R, or KNull to interact with DnaJ-1 (figure 4C). Lastly, mRNA levels of DnaJ-1 were not significantly impacted by the various forms of Atxn3 expressed in this study (figure 4D). Due to lack of reliable antibodies to detect endogenous DnaJ-1 protein in the fly, we were not able to examine this co-chaperone at the protein level. We conclude that lysine mutations do not impact the protective role that DnaJ-1 has on the toxicity of pathogenic Atxn3.

**Figure 4:**
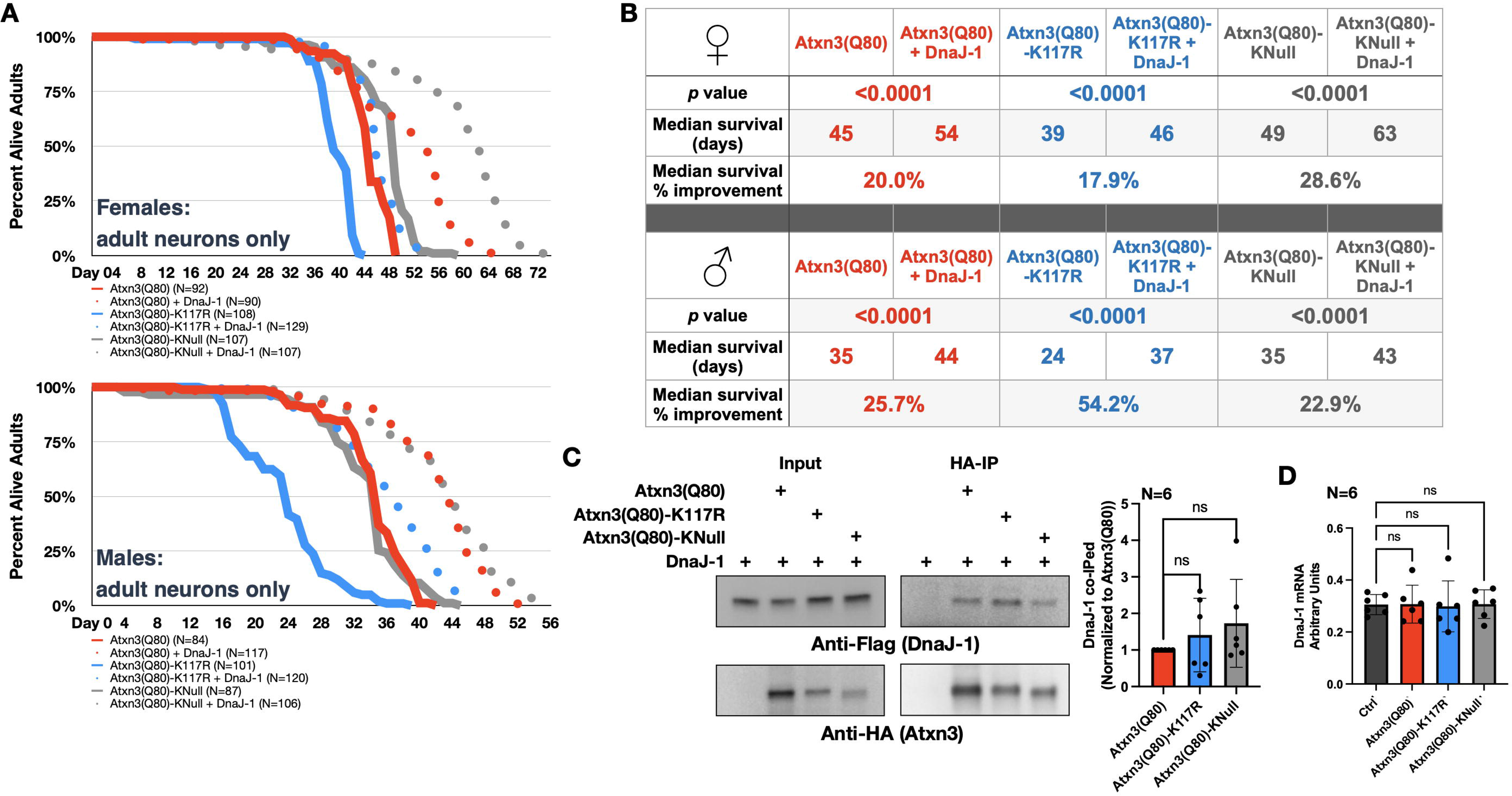
DnaJ-1 suppresses toxicity of pathogenic Atxn3 independently of lysine mutations. A, B) Longevity outcomes from adult flies expressing the noted versions of Atxn3 in the presence or absence of exogenous DnaJ-1, only in adult neurons. B) Statistics: log-rank tests. C) Left: Western blots from co-immunopurifications of the Atxn3 versions noted expressed in female adult neurons for 7 days in the presence of exogenous FLAG-tagged DnaJ-1. Right: quantifications from the left and other independent repeats. Statistics: Wilcoxon tests. ns: not significant. FLAG-tagged DnaJ-1 was used for these studies since antibodies to detect endogenous DnaJ-1 are not available. D) qRT-PCR of endogenous levels of DnaJ-1 when Atxn3 of the noted versions was expressed. Statistics: one-way ANOVA with Dunnett’s post-hoc. ns: not significant.

### K117 mutation modestly increases the aggregation of pathogenic Atxn3

Thus far, we have observed mild but significant differences in the toxicity Atxn3(Q80) versus K117R, but have not identified clues that elucidate a mechanism underlying these observations: overall protein levels are the same; their turnover is comparable; they are distributed similarly between cytoplasm and nucleus; and exogenous DnaJ-1 is able to successfully rescue all versions. We next turned our attention to the aggregation of Atxn3 in fly neurons. In our prior work, we found that aggregation of pathogenic Atxn3 correlates with its toxicity, whether measured by changes in longevity, motility, or tissue structure (Johnson et al., 2021; Johnson et al., 2020; Prifti et al., 2022; Ristic et al., 2018; Sutton et al., 2017; Tsou et al., 2015b).

We expressed Atxn3(Q80), K117R, or KNull in adult fly neurons for 1, 7 or 14 days. We then resolved the expressed protein through SDS-PAGE gels and quantified SDS-soluble versus SDS-resistant species for each lane. As summarized in figure 5, we noticed a modestly higher preponderance of SDS-resistant species with K117R compared to Atxn3(Q80) as time progressed, especially in female flies; male flies showed a trend in day 14, but did not reach significance. For calculations, we quantified the amount of SDS-soluble and SDS-resistant protein in each lane and graphed the soluble portion as a fraction of the total amount of Atxn3 signal in the respective lane. Based on these assays on days 7 and 14, KNull was consistently more aggregation-prone than either of the other two versions of pathogenic Atxn3, even though the total protein levels of KNull were lower than those of Atxn3(Q80) and K117R, as also shown in figure 3. These data from adult neurons suggest a small trend of increased aggregation of Atxn3(Q80) when K117 or all of its lysines are mutated into arginine residues (figure 6).

**Figure 5:**
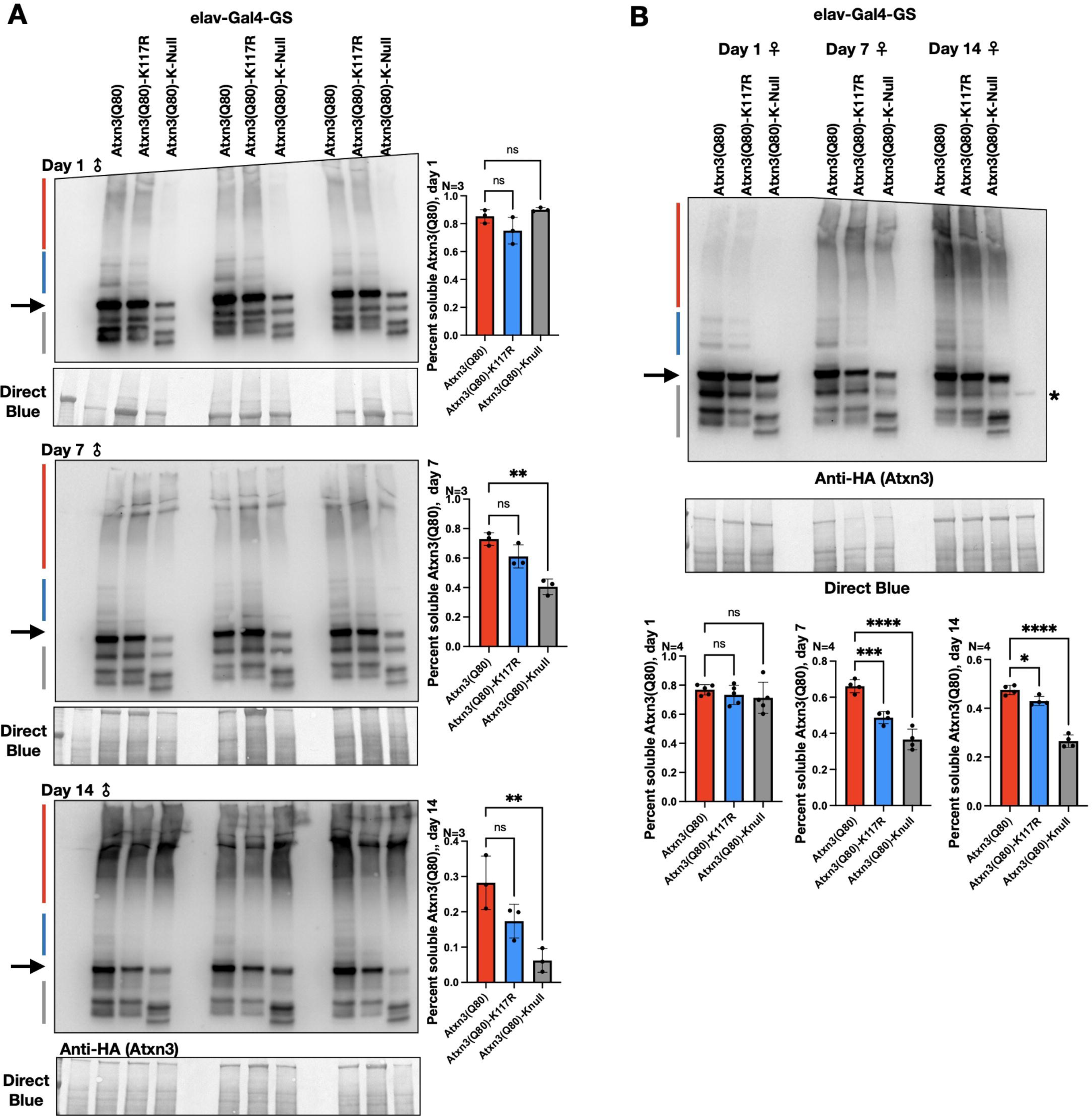
Lysine mutations increase the aggregation propensity of pathogenic Atxn3 in adult neurons. A, B) Western blots from adult flies expressing the noted versions of Atxn3 in adult neurons for the indicated amounts of time. Black arrow: main, unmodified Atxn3 band. Blue arrows: ubiquitinated Atxn3. Red bar: SDS-resistant Atxn3. Gray bar: proteolytic fragments of Atxn3. Associated quantifications are from blots; percent soluble denotes “non SDS-resistant.” Means -/+ SD. Atxn 3 signal for each lane was calculated by quantifying SDS-soluble species, SDS-resistant species, and expressed as percent soluble fraction of the total Atxn3 signal in that lane. Statistics: one-way ANOVA with Dunnett’s post hoc. ns: not significant. *: p<0.05. ***: p<0.001. ****: p<0.0001.

**Figure 6:**
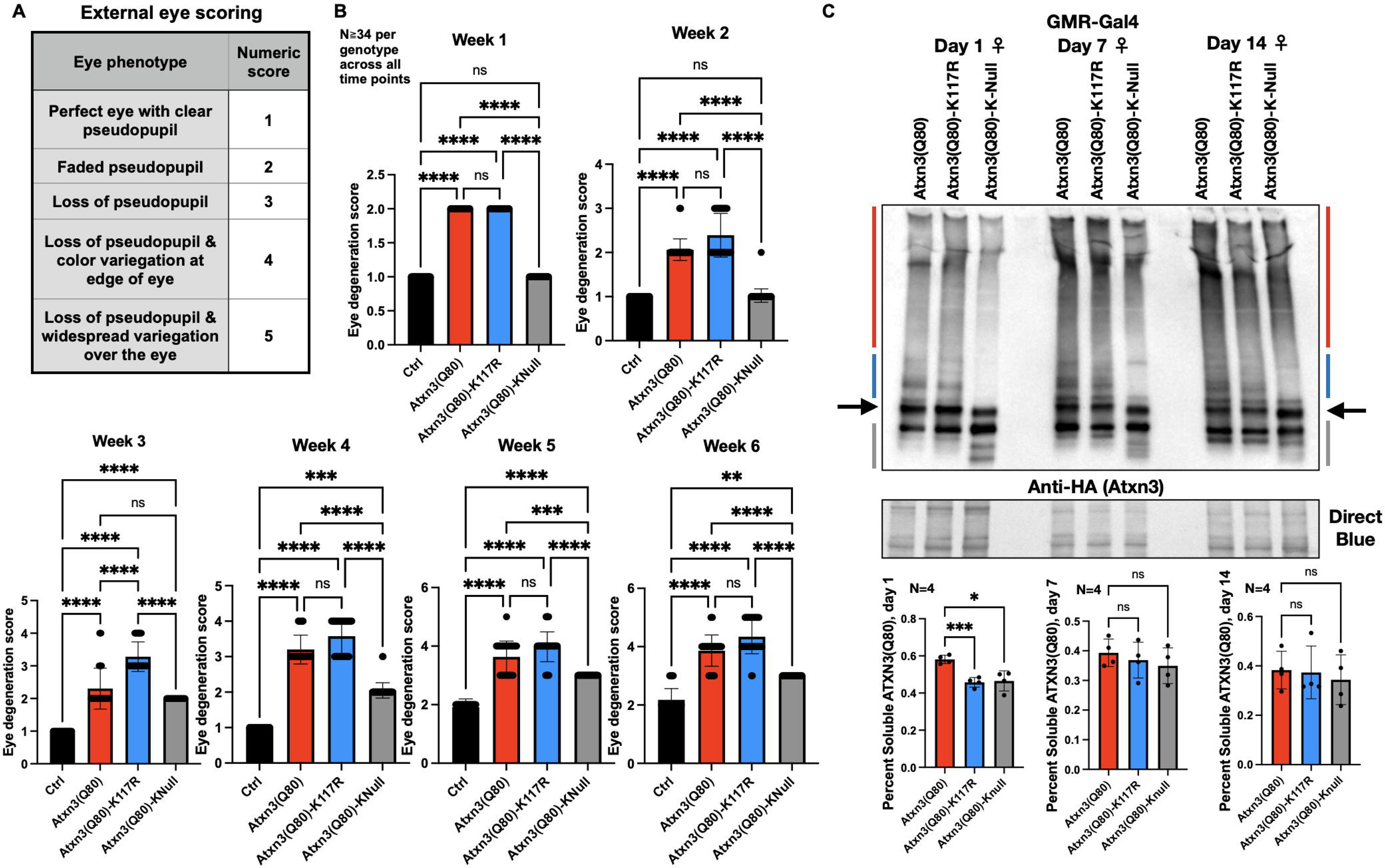
Lysine mutations impact the toxicity of pathogenic Atxn3 in fly eyes. A) Eye scoring system. B) Means -/+ SD of eye phenotypes when the noted versions of Atxn3 are expressed in fly eyes (GMR-Gal4 driver). ns: not significant. **: p<0.01. ***: p<0.001. ****: p<0.0001. C) Western blots and related quantification from adult female fly heads, day 1, homogenized to examine levels of the Atxn3 versions noted. Blue arrows: ubiquitinated Atxn3. Red bar: SDS-resistant Atxn3. Gray bar: proteolytic fragments of Atxn3. Quantifications are from top and additional independent repeats. Atxn 3 signal for each lane was calculated by quantifying SDS-soluble species, SDS-resistant species, and expressed as percent soluble fraction of the total Atxn3 signal in that lane. Means -/+ SDS. Statistics: Brown-Forsythe and Welch ANOVA tests. ns: not significant; *: p<0.05. ***: p<0.001.

To enhance the overall rigor of our work, we additionally tested the toxicity of pathogenic Atxn3 versions in fly eyes. Using a scoring mechanism that we reported in the past (Johnson et al., 2020; Prifti et al., 2022) (figure 7A), we noted that differences among Atxn3(Q80), K117R and KNull that we observed in other tissues are recapitulated in fly eyes: K117 mutation worsens the phenotype observed with Atxn3(Q80), whereas KNull mutations improve them.

These differences are statistically significant on day 21 for Atxn3(Q80) versus Atxn3(Q80)- K117R, and are significant between Atxn3(Q80) and Atxn3(Q80)-KNull at all points examined. At the biochemical level, we observed increased aggregation propensity with Atxn3(Q80)- K117R and KNull on day 1 compared with Atxn3(Q80), but not at later time points. These collective results suggest that mutating either K117 or all lysine residues on pathogenic Atxn3 can increase its tendency to aggregate in fly tissue; but, this is a modest effect.

## DISCUSSION

Atxn3, whose polyQ tract expansion causes SCA3, is a DUB. Its enzymatic activity is enhanced by ubiquitination of K117. Here, we asked: is K117 important for the toxicity of pathogenic Atxn3? We found that the answer is yes, but not too much. This answer rests on our data that expression of pathogenic Atxn3 with a mutated K117 into arginine was more toxic than polyQ-expanded Atxn3 with an intact K117 when expressed in various fly tissues and at different stages of the fly life cycle. This mildly enhanced toxicity did not coincide with clear differences in the total protein levels, its sub-cellular distribution, or its interaction with the co-chaperone DnaJ- 1, but correlated with a small trend of increased aggregation of the K117R version. Our observational results do not point to a mechanistic understanding of how the K117R mutation enhances toxicity, but we presume that it results from reduced catalytic activity of Atxn3, impeding its functions. We will pursue this possibility as further details about the precise functions of Atxn3 in vivo continue emerging.

In addition to K117R, we also generated a version of pathogenic Atxn3 whose individual lysine residues are mutated into arginines, denoted as KNull. This transgene was generated to ask the question: 1) is ubiquitination of pathogenic Atxn3 necessary for its degradation in vivo? We had previously documented that wild-type Atxn3 does not need to be ubiquitinated to be degraded in mammalian cells or in intact flies (Blount et al., 2020; Blount et al., 2014), and that pathogenic Atxn3 also does not need to be ubiquitinated to be degraded in mammalian cells (Blount et al., 2020); still the question remained whether ubiquitination is essential for the degradation of pathogenic Atxn3 in vivo. 2) We also wondered if overall ubiquitination impacts pathogenic Atxn3 toxicity. The answer to the first question confirms our previous reports that ubiquitination is not an absolute requirement for the turnover of pathogenic Atxn3. This is not to say that there are not instances when ubiquitination regulates Atxn3 degradation; prior evidence towards this role is robust and instances when ubiquitination fine tunes the degradation of this DUB are highly likely (Dantuma and Herzog, 2020; Johnson et al., 2022b; Lieberman et al., 2018; Matos et al., 2019b). The answer to the second question is not as straightforward. According to biochemical examinations, KNull Atxn3(Q80) is more aggregation prone than the version with all lysines present. However, the KNull version is also less toxic. We propose that reduced toxicity is due to the markedly lower levels of Atxn3(Q80)-KNull protein, potentially resulting from translation-related degradation, which is a point that needs future investigation. We conclude that lysine residues are important for both the aggregation and toxicity of pathogenic Atxn3. Additional studies are needed to dissect the contribution of individual lysine residues in these processes.

We noticed a likely correlation between increased aggregation and toxicity of pathogenic Atxn3 in flies as a result of lysine mutations. Generally, we observed modestly increased trends of aggregation when K117 or all lysines were mutated on pathogenic Atxn3; these trends reached significance at some time points and led us to propose that the aggregative propensity of pathogenic Atxn3 is in part regulated by lysine residues. For example, ubiquitinated Atxn3 may be less aggregation-prone due to steric hindrances that oppose oligomerization of the disease protein. An additional, non-mutually exclusive mechanism may rely on ubiquitinated Atxn3 partnering up with additional proteins through ubiquitin-binding sites, which can also hinder aggregation. There is prior evidence from the polyQ field that ubiquitination of a disease protein impacts its aggregation and toxicity, as reported for the Kennedy Disease protein, androgen receptor (Pluciennik et al., 2021; Sengupta et al., 2022).

Various ubiquitin-related enzymes have been reported to ubiquitinate Atxn3, including the E2 conjugases UbcH5C and Ube2W, and the E3 ligases CHIP and Parkin. In our prior work (Tsou et al., 2013) and in additional experiments we conducted for this study (Supplemental figure 1), we have not observed differences when Ubch5C or CHIP were knocked down in flies. We propose that Atxn3 is modified in vivo by more than one specific conjugase/ligase pair, and that individual targeting is insufficient to modify its ubiquitination pattern in this system.

The precise functions of Atxn3 remain to be elucidated. Studies conducted with reconstituted systems in vitro, in cultured mammalian cells, and in mouse models reveal a DUB that is involved in some aspects of protein quality control, including ER-Associated Degradation and autophagy (Liu and Ye, 2012; Zhong and Pittman, 2006); an enzyme that functions in sensing the overall levels of ubiquitin in the system (Dantuma and Herzog, 2020; Scaglione et al., 2011; Winborn et al., 2008); that regulates DNA-damage repair processes (Dantuma and Herzog, 2020; Matos et al., 2019a); and whose wild-type protein is involved in the visual system (Toulis et al., 2020). As a gene, *Atxn3* appears to be dispensable in mice (Reina et al., 2010; Schmitt et al., 2007; Switonski et al., 2011), the pathogenic polyQ expansion likely perturbs some of the functions of wild-type Atxn3 and likely imparts news ones onto the SCA3 protein, resulting in neuronal and glial malfunction, clinical manifestation and neurodegeneration.

Ubiquitination of Atxn3 at K117 markedly enhances its activity as an enzyme in vitro and upregulates some of its cellular functions, such as aggresome formation and protections against proteotoxic stress (Burnett and Pittman, 2005; Todi et al., 2010; Tsou et al., 2013).

Ubiquitination of pathogenic Atxn3 at K117 may enhance some of its normal functions, enhance some of its toxic, gain-of-function properties, or both. Based on our results that prohibiting ubiquitination at K117 increases the toxicity of pathogenic Atxn3 in flies, we suggest that ubiquitination of K117 on Atxn3 is largely protective in SCA3. The exact details require further investigation to understand the biology of disease of SCA3.

## METHODS

### Antibodies

Anti-HA (rabbit monoclonal C29F4, 1:500-1000; Cell Signaling Technology) Anti ataxin-3 (mouse monoclonal 1H9, MAB5360, 1:500-1000; Millipore)

Anti-lamin (mouse monoclonal ADL84.12-5, 1:1000; Developmental Studies Hybridoma Bank) Anti-tubulin (mouse monoclonal T5168, 1:10,000; Sigma-Aldrich)

Anti-Flag (rabbit polyclonal SAB4301135, 1-1000; Sigma-Aldrich)

### Drosophila lines

Atxn3 transgenic lines were generated as described before. Briefly, human Atxn3 cDNA was sub-cloned into pWalium 10.moe vector for PhiC31 integration into site AttP2 of the third chromosome of the fly. Transformants were examined by PCR for integration site and direction, and sequenced for construct integrity. sqh-Gal4 was a gift from Dr. Daniel Kiehart, Duke University. elav-Gal4 was a gift from Dr. Daniel Eberl, University of Iowa, and elav-Gal4- GS was a gift from Dr. R. J. Wessells, Wayne State University. GMR-Gal4 was stock #8121 from Bloomington *Drosophila* Stock Center. Control flies consisted of a single outcross of the respective Gal4 driver into the same genetic background as each Atxn3 transgenic line, without expressing Atxn3.

### *Drosophila* husbandry and longevity

Flies were maintained at 25°C, diurnally controlled incubators with 12 h light/dark cycles and on conventional cornmeal media. All offspring were maintained in the same conditions, with the exception of crosses that utilized elav-GS-Gal4. Those offspring were switched into media containing RU486 soon after emerging as adults to induce the expression of transgenes. For lifespan recordings, adult flies were collected each day onto 10% sucrose 10% yeast (SY10) media and separated into groups of the same size (10 adults per vial) and gender. Flies were flipped into new SY10 food every other day. The number of adults tracked (N) for each group is noted in figures. In case of developmental lethality, flies were observed from the embryo stage through eclosure or early adult death, and lethality was recorded multiple times per week.

### Western blotting

Western blotting was conducted using 5 adult flies (neuronal expression) or 5 dissected fly heads (eye expression) per sample. Samples were homogenized in boiling SDS lysis buffer [50 mM Tris pH 6.8, 2% SDS, 10% glycerol, 100 mM dithiothreitol (DTT)], sonicated, boiled for 10 min, and subsequently centrifuged for 10 min at 13,300 rpm at room temperature. Samples were electrophoresed through Tris/Glycine gels (Bio-Rad). Western blots were imaged using ChemiDoc (Bio- Rad). Quantification was conducted using ImageLab (Bio-Rad). We used direct blue stains of total protein as loading controls by saturating PVDF membrane for 10 min in 0.008% Direct Blue 71 (Sigma-Aldrich) dissolved in 40% ethanol and 10% acetic acid, rinsed with a solution of 40% ethanol/10% acetic acid, then ultra-pure water, and then air dried and imaged.

### Co-immunopurifications

Ten adult flies expressing specific transgenes through the elav-Gal4- GS driver were thoroughly homogenized in 800 µL of buffer consisting of NETN or 1:1 NETN/PBS supplemented with protease inhibitors (PI; Sigma-Aldrich, S-8820). Homogenates were rotated at 4°C for 25 minutes, and then centrifuged for 10 minutes at 5000 x g at 4°C. Supernatant was subsequently combined with bead-bound antibody and tumbled at 4°C for 4 hours. Beads were rinsed 5 times each with lysis buffer, and bead-bound complexes were eluted through Laemmli buffer (Bio-Rad) and heating at 95°C for 5 minutes. Supernatant was then supplemented with 6% SDS to a final concentration of 1.5% SDS and loaded for Western blotting.

### qRT-PCR

Relative message abundance from 6 individual samples per genotype was determined by amplification and staining with Power SYBR Green using the QuantStudio3 system and Design and Analysis 2.6.0 software (Applied Biosystems). *Rp49* abundance was used to calculate relative expression. Primer sequences are listed below:

rp49-F: 5′-AGATCGTGAAGAAGCGCACCAAG-3′, rp49-R: 5′-CACCAGGAACTTCTTGAATCCGG-3′ ataxin-3-F: 5′-GAATGGCAGAAGGAGGAGTTACTA-3′ ataxin-3-R: 5′-GACCCGTCAAGAGAGAATTCAAGT-3′ DnaJ-1-F1: 5′-GTACAAGGAGGAGAAGGTGCTG-3′ DnaJ-1-R1: 5′-CAGACTGATCTGGGCTGTATACTT-3′

### Nuclear/cytoplasmic fractionation

Fractionation of nuclear and cytoplasmic fractions from *Drosophila* lysates was conducted with the ReadyPrep Protein Extraction Kit (Bio-Rad). Seven whole flies per group were homogenized in cytoplasmic extraction buffer (Bio-Rad) and subsequently processed as instructed by the manufacturer.

### Statistics

Statistical analyses were conducted with GraphPad’s Prism 9. Tests used are specified in the figure legends.

## ACKNOWLEDGEMENTS

We extend our gratitude to Era Cobani for her help with conducting several of the lifespan assays.

## AUTHOR CONTRIBUTIONS

JRB: conceptualization, data curation, software, formal analysis, validation, investigation, visualization, methodology, and editing.

NCP: data curation, software, formal analysis, validation, investigation, visualization, methodology, and editing.

KL: data curation, validation, investigation, methodology.

ALH: data curation, funding acquisition, validation, investigation, methodology.

W-LT: conceptualization, data curation, software, formal analysis, validation, investigation, visualization, methodology.

AS: conceptualization, data curation, software, formal analysis, validation, investigation, visualization, methodology, and writing and editing.

SVT: conceptualization, data curation, funding acquisition, software, formal analysis, validation, investigation, visualization, methodology, and writing and editing.

## SUPPLEMENTAL FIGURES

**Supplemental figure 1.** Outcomes from crosses of flies expressing the noted versions of Atxn3 in fly eyes (GMR-Gal4) alongside RNAi targeting the noted genes. Dissected and homogenized heads for the Western blots were from 1-day-old female flies. Each lane is an independent repeat. Black arrow: main, unmodified Atxn3 band. Blue bar: ubiquitinated Atxn3. Red bar: SDS-resistant Atxn3. Gray bar: proteolytic fragments of Atxn3. RNAi and over-expression (O/E) stocks were as follows: Ubch5C: RNAi-1, BDSC 31875; RNAi-2, BDSC 35431; O/E: BDSC 26691 for HA-tagged, and BDSC 56495 for untagged. CHIP: RNAi-1, BDSC 33938; RNAi-2: 34017; O/E:

