## Supplementary figures and images for "Lysine 117 on ataxin-3 modulates toxicity in *Drosophila* models of Spinocerebellar Ataxia Type 3"

### Supplemental File 1

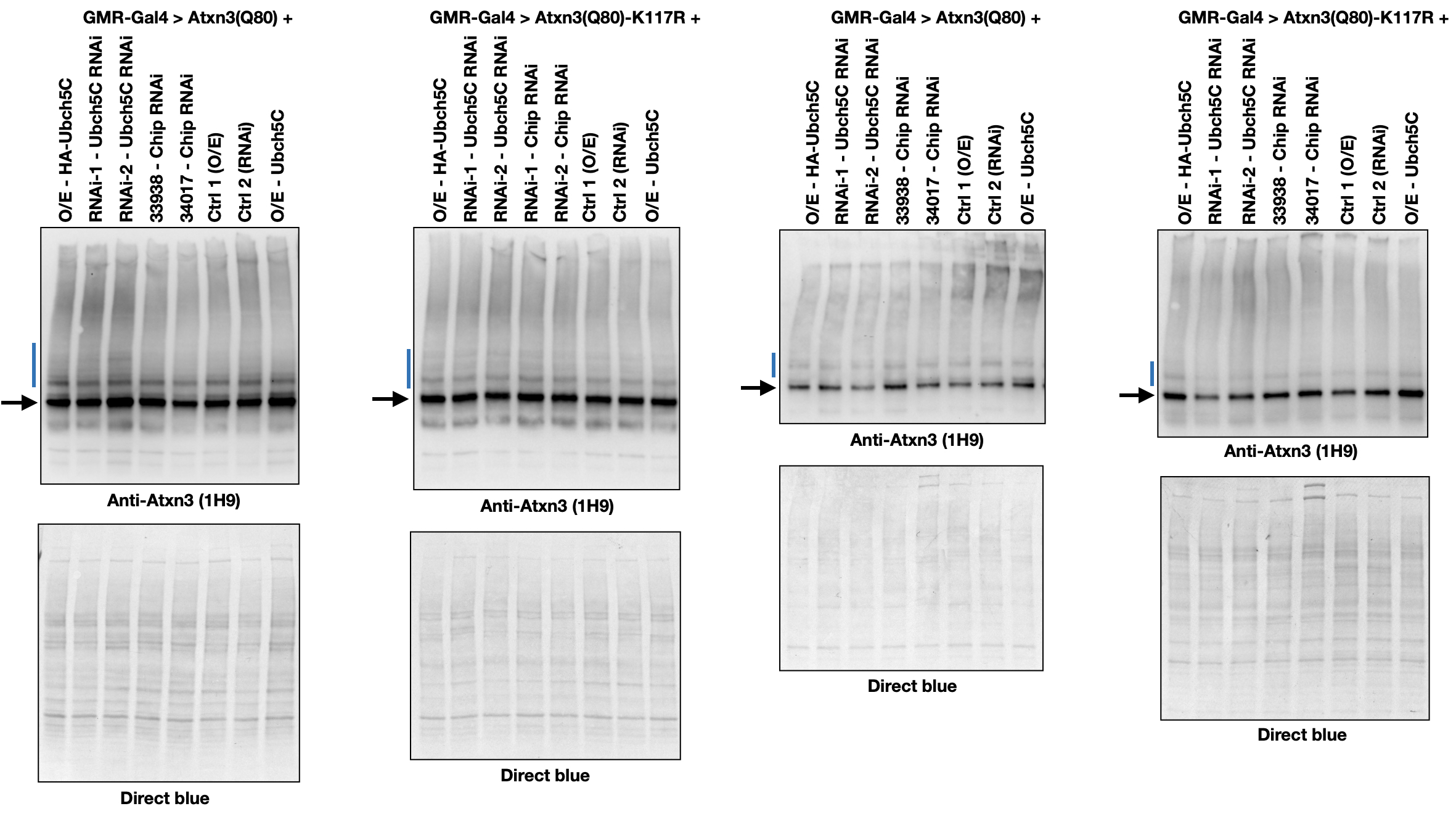
